# Efficient rescue of a newly classified Ebinur lake orthobunyavirus with GFP reporter and its application in rapid antiviral screening

**DOI:** 10.1101/2022.03.25.485793

**Authors:** Nanjie Ren, Fei Wang, Lu Zhao, Shunlong Wang, Guilin Zhang, Jiaqi Li, Bo Zhang, Eric Bergeron, Zhiming Yuan, Han Xia

## Abstract

Orthobunyaviruses have been reported to cause severe diseases in humans or animals, posing a threat to human health and social economy. Ebinur lake virus (EBIV) is a newly classified orthobunyavirus, which needs further intensive study and therapies to cope with its potential infection risk to human and animals. Here, through the reverse genetics system, the recombinant EBIV of wild type (rEBIV/WT) and NP-conjugated-eGFP (rEBIV/eGFP/S) were rescued for the application of the rapid antiviral drug screening. The eGFP fluorescence signal of the rEBIV/eGFP/S was stable in the process of successive passage in BHK-21 cells (over 10 passages) and this recombinant virus could replicate in various cell lines. Compared to the wild type EBIV, the rEBIV/eGFP/S caused the smaller plaques and its peak titers were lower, suggesting attenuation due to the eGFP insertion. Through the high-content screening (HCS) system, ribavirin showed an inhibitory effect on the rEBIV/eGFP/S with an EC50 of 21.91 μM, while favipiravir did not inhibit, even at high concentrations. In addition, five of ninety-six natural compounds had antiviral against EBIV. The robust reverse genetics system for EBIV will facilitate investigation into replication and assembly mechanisms and assist drug and vaccine development.

## 1 Introduction

Emerging and re-emerging arboviruses pose a big threat to human and animal health worldwide (1). The genus *Orthobunyavirus* (family *Peribunyaviridae*) includes over 18 serogroups is the largest genus in the order *Bunyavirales*, comprising over 170 arboviruses which were mainly transmitted through mosquito vectors (2)(3). These negative-sense RNA viruses take their name from Bunyamwera virus (BUNV), which was originally isolated in 1943 from *Aedes* mosquitoes during an investigation of yellow fever in the Semliki Forest, Uganda (4). Some members of *Orthobunyavirus* can result in several disease syndromes in humans, including acute but self-limiting febrile illness (Oropouche virus (OROV)) (5), pediatric arboviral encephalitis (La Crosse virus (LACV)) (6) and haemorrhagic fever (Ngari virus (NRIV)) (7). Apart from human pathogens, orthobunyaviruses comprises some veterinary pathogens, such as Akabane virus (AKAV) and Schmallenberg virus (SBV), which both cause congenital malformations in ruminants (8)(9).

Orthobunyaviruses are tri-segmented negative-sense RNA viruses, comprising small (S), medium (M), and large (L) segments (4), named by the segment length. The *L* segment encodes the viral RNA-dependent RNA polymerase (RdRp), which involved in the genome replication. The *M* segment encodes two structural glycoproteins, Gn and Gc, and a non-structural protein (NSm). Gn and Gc are related to receptor binding and the fusion of the viral and endosomal membranes (10). The NSm was suggested to function as a scaffold for virion assembly (11). The *S* segment encodes the nucleocapsid protein (N) and a non-structural protein (NSs) in overlapping open reading frames with the same direction. The N protein encapsidates both genomic and antigenomic RNA (but not viral mRNA) to form ribonucleoprotein (RNPs) complexes that are the templates for the viral RdRp (4)(12). The NSs protein is considered as the major virulence determinant of orthobuyaviruses by antagonizing host innate immune responses, including type I interferon responses (13)(14).

Reverse genetic systems are powerful and versatile molecular tools for the study of RNA viruses, which can be used to produce attenuated virus vaccine (15), probe viral replication and interactions with host innate immune responses (4). Since the BUNV was recovered from transfecting cells with just three plasmids that express full-length antigenome viral RNAs in 2004 (16), the infectious cDNA clones have been obtained for several orthobunyaviruses, such as SBV (17), LACV (18), AKBV (19) and Shuni virus (SHNV) (10). According to the previous researches, we can find that two transcription plasmids for the reverse systems have been developed: (1) transcription plasmids based on T7 promoter which could be recognized by T7 RNA polymerase; (2) transcription plasmids using mammalian RNA polymerase I promoters (4). Considering the efficiency of RNA polymerase I-driven system was lower than that of T7 RNA polymerase (19), the latter was chosen in our study.

Ebinur lake virus (EBIV) is isolated from *Culex modestus* mosquito pools in Ebinur lake region, Xinjiang in 2014 (20). Our previous work demonstrates that EBIV has the highest similarity with Germiston virus (GERV) (21), which belongs to risk group 3 human pathogens. Remarkably, EBIV can cause acute lethal disease in adult mice (22), and the antibodies against EBIV are detected in local residents, which indicate that EBIV has a potential infection risk in animal and/or human (21). Hence, we use the reverse genetics system to rescue the recombinant EBIV of wild type (rEBIV/WT) and NP-conjugated-eGFP (rEBIV/eGFP/S). Furthermore, through the high-content screening (HCS) system, ribavirin and five natural compounds was found to have the antiviral effects against the recombinant EBIV.

## 2 Methods and Materials

### 2.1 Cells, viruses and antibodies

Baby hamster kidney cell (BHK-21), African green monkey kidney cells (Vero E6), human adrenal cortical carcinoma cells (SW13), and porcine kidney epithelial cells (PK15) were propagated in Dulbecco’s modified Eagle’s medium (DMEM, Gibco) supplemented with 10% fetal bovine serum (FBS, Gibco), 100 units/mL of penicillin and 100 μg/mL of streptomycin. BSR-T7 cell, a generous gift from Prof. Bo Zhang, which could express T7 RNA polymerase, were cultured in DMEM supplemented with 10% FBS and 1 mg/mL G418 (Beyotime). All mammalian cells were grown at 37 °C under a 5% CO_2_ atmosphere. The chicken hepatocellular carcinoma cell line (LMH) was maintained in Dulbecco’s Modified Eagle Medium/Nutrient Mixture F-12 (DME/F-12, HyClone) containing 10% FBS and 1% penicillin/streptomycin at 37 °C in 5% CO_2_. The Aedes albopictus mosquito cell (C6/36) was grown in Roswell Park Memorial Institute (RPMI) 1640 medium supplemented with 10% FBS and 1% penicillin/streptomycin, and were maintained in 5% CO_2_ at 28 °C.

EBIV isolate Cu20-XJ was first isolated from *Culex modestus* mosquitoes in Xinjiang, China (20). The EBIV virus stock was propagated in BHK-21 cells in DMEM containing 2% FBS, subpackaged and stored at −80 °C. The rEBIV/WT and reporter virus rEBIV/eGFP/S were produced through the transfection of the plasmids (described below) into BSR-T7 cells with transcription plasmids (described below). All the work with infectious virus were conducted in the biosafety level-2 (BSL-2) laboratory.

The mouse polyclonal antibody against EBIV N protein was generated by immunization of BALB/C mice with the purified EBIV N protein. Alexa Fluor™ 594 goat anti-mouse IgG (H+L) used as secondary antibody was purchased from Invitrogen.

### 2.2 Sequencing of EBIV genome 5’ ends

EBIV were first concentrated by adding 3.2 g polyethylene glycol (PEG) 8000 (8%, W/V, MilliporePEG8000, Sigma, USA) and 0.9 g NaCl (0.3 mol/L, Millipore Sigma, USA) to 40 mL virus supernatant (23). The suspension was mixed vigorously and incubated overnight at 4°C. Then the supernatant was discarded after centrifugation at 9000 × g for 30 min at 4°C. The pellet was resuspended in 200 μL PBS. Then the viral RNA was isolated using Trizol reagent (Invitrogen). First strand cDNA synthesis was carried out with one microgram RNA, Random Primer Mix (Takara) and 200 U SMARTScribe Reverse Transcriptase (Takara). EBIV M/L segment-specific sequences were amplified by PCR including Universal Primer A Mix (Takara), Gene-Specific Primers (GSPs) (M: 5’-GATTACGCCAAGCTTAGAACTAGTAGGTGGGGCTGCGAAG-3’; L: 5’-GATTACGCCAAGCTTGGACTAAGATGTTGACGCAGCAGGAT-3’), 2.5 µl cDNA and 1.25 U SeqAmp™ DNA polymerase (Takara). The PCR products were separated in a 1% agarose gel and recovered with a Gel extraction kit (Omega). While the PCR products were weak or smear bands, the PCR would be conducted again with diluted PCR products, Universal Primer Short and Nest Gene-Specific Primers (NGSPs) (M: 5’-GATTACGCCAAGCTTGTGCCATATCAGGACCCTGTGAGACC-3’; L: 5’-GATTACGCCAAGCTTGGAGGAGAAATGAGGAAGGCAATC-3’). Amplicons were cloned into vector pRACE (Takara) and individual clones were selected for nucleotide sequencing.

### 2.3 Plasmid construction

Full-length cDNA from the S, M, and L segments (GenBank accession no. KJ710423, KJ710424, KJ710425) was obtained by reverse transcription of EBIV RNA using GoScript™ Reverse Transcriptase (Promega, USA) and a pair of DNA primers complementary to the 5’ and 3’ end of the viral genomic RNAs. Complete cDNAs were sequence-amplified by PCR using KOD One™ PCR Master Mix -Blue- (TOYOBO, Japan) and were cloned into pSMART-LCK plasmid (Lucigen, Middleton, WI, USA) containing a T7 RNA polymerase promoter, hepatitis D ribozyme and T7 RNA polymerase terminator motif (abbreviated to pLCK), (24). As shown in Fig. 1A, C and D, the resulting plasmids (pLCK-EBIV-S, pLCK-EBIV-M, and pLCK-EBIV-L) containing viral different segment sequences located between a T7 promoter and a hepatitis D ribozyme T7 polymerase terminator motif. In the plasmid pLCK-EBIV-eGFP/S, the *eGFP* gene was fused with the porcine teshovirus-1 2A peptide linker sequence (P2A), a together inserted before the ORF of EBIV S segment (Fig. 1B). The P2A peptide is a self-cleaving peptide allowing separate expression of two proteins via a ribosomal skipping event during it translation (25). The reporter virus rEBIV/eGFP/S was generated through the transfection of the plasmids (pLCK-EBIV-eGFP/S, pLCK-EBIV-M, and pLCK-EBIV-L) into BSR-T7 cells. All the constructs were confirmed by sequencing and submitted to the Genbank (Acession No: ON055165 to ON055168).

**Fig 1.**
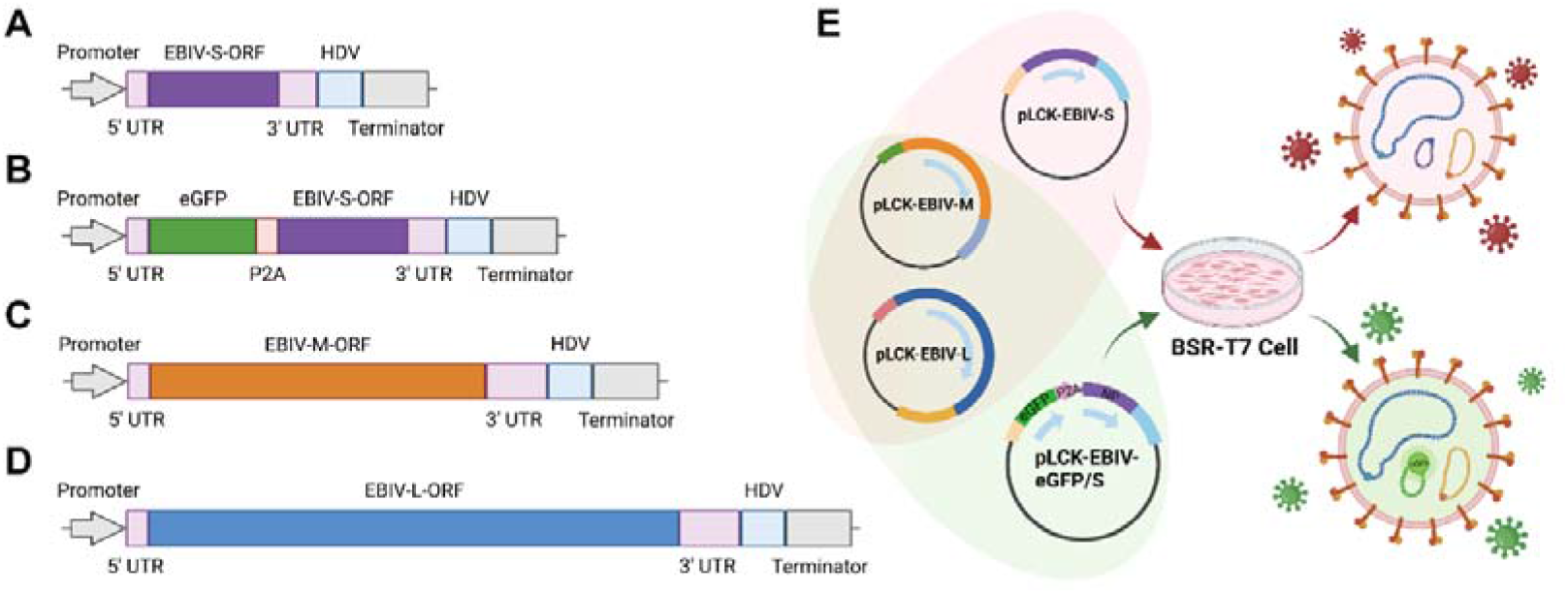
Structure diagram and transfection strategy of the recombinant plasmids rescue the recombinant EBIV. (A) The recombinant pLCK plasmid of S segment, named pLCK-EBIV-S. (B) Based on pLCK-EBIV-S, eGFP and P2A gene were added between 5’ UTR and N protein ORF, named pLCK-EBIV-eGFP/S. (C) The recombinant plasmid of M segment, named pLCK-EBIV-M. (D) The recombinant pLCK plasmid of L segment named pLCK-EBIV-L. (E) Transfection strategy for rEBIV/WT and rEBIV/eGFP/S. Based on the pSMART-LCK plasmid, the pLCK plasmid were reconstructed by inserting the T7 promoter upstream the 5’ UTR and HDV downstream the 3’ UTR.

### 2.4 Virus rescue and plaque assay

Rescue of recombinant viruses was performed in BSR-T7 cells, which constitutively express T7 polymerase. As shown in Fig 1E, a 50% confluent monolayer of BSR-T7 cells grown in 12-well plates was transfected with 300 ng pLCK-EBIV-S, 500 ng pLCK-EBIV-M, and 700 ng pLCK-EBIV-L. When obvious cytopathic effect (CPE) appeared, supernatants contained the rEBIV/WT were harvested.

For the rescue of reporter virus, the pLCK-EBIV-eGFP/S (500ng), which was the substitute of pLCK-EBIV-S, was added in the transfection. At the next day post transfection, the green fluorescence could be observed by the inverted fluorescent microscope. The supernatants of cell culture were harvested at 120 h post transfection, labeled as rEBIV/eGFP/S and stored at −80 °C.

The rescued viruses were subjected to a serial dilution of 10-fold with DMEM until 10^−6^. Add 100 µl virus diluent into each well of 24-well plates containing the monolayer BHK-21. After 1 h incubation, the virus diluent was discarded, and each well was were added with 500 µl DMEM covering containing 1.5% methyl cellulose. The cells would be cultured at 37 °C in 5% CO_2_. Three days later, the cells were fixed overnight with 3.7% formaldehyde and stained with 2% crystal violet for 15 min. The amount and size of plaques were recorded.

### 2.5 Immunofluorescence assay

The EBIV-WT, rEBIV/WT and rEBIV/eGFP/S viruses were seeded on a 12-well plate containing BHK-21 cell, respectively. After 24 hours, the cells were fixed in cold (−20 °C) 100% methanol for 5 min at −20 °C and washed three times with PBS. For the permeabilization, add the PBS containing 0.1% Triton X-100 and incubate them for 15 min at room temperature. After that, wash them with PBS three times, add the PBS containing 2% BSA, and incubated them for 60 min at room temperature. Then the cells were incubated with the mouse polyclonal antibody against EBIV-NP (1:250 diluted in PBS containing 0.1% BSA) overnight at 4°C. After washing with PBS three times, the cells were incubated with goat anti-mouse IgG conjugated with Alexa Fluor 594 (1:250 diluted in PBS with 0.1% BSA) at room temperature for 1 h (avoid light). Following three times of PBS washing, the DAPI was added to the cells, and keep in for 5 min at room temperature. Then the fluorescence signal of each well was observed and analyzed under an Olympus fluorescence microscope at 200 × magnification (26).

### 2.6 Viral growth kinetics

To compare the differences in replication among the three viruses, the growth kinetics of EBIV-WT, rEBIV/WT and rEBIV/eGFP/S viruses on BHK-21 were examined respectively. Approximately 1 × 10^4^ BHK-21 cells were seeded in a 17.5 mm dish. After incubation overnight, the cells were infected with 1mL EBIV-WT, rEBIV/WT or rEBIV/eGFP/S virus at MOI of 0.01. After incubation for one hours, the supernatants were collected, the cells were washed with PBS for three times and replaced with fresh medium with 2% FBS. Every 24 hours post infection, the supernatants were collected and stored at −80 °C. At last, they were subjected to plaque assay to determine the viral titer. For rEBIV/eGFP/S infection, the expression of eGFP gene was observed under the fluorescence microscope at 100 × magnification.

### 2.7 Stability of the rEBIV/eGFP/S virus in cell culture

To analyze whether the eGFP reporter gene can be stable presence during the passage, the rEBIV/eGFP/S virus was serially passaged in vertebrate-derived BHK-21 cells for ten rounds respectively. For each generation, 200 μL viruses were used to infect naïve BHK-21 cells, and the percentage of cells expressing eGFP was evaluated at 24-48 hpi (hour post infection) after each passage. In addition, the RNAs of the infected cells were extracted in each passage and separated into two parts, one subjected to RT-PCR using PrimeScript™ One Step RT-PCR Kit Ver.2 (Dye Plus) (Takara, Japan), then the region between S-5’UTR and N protein-ORF genes was amplified to monitor the expression of eGFP.

The rest of RNAs were performed with real-time reverse transcription PCR (RT-qPCR) using a Luna^®^ Universal Probe One-Step RT-qPCR Kit (New England Biolabs) according to the manufacturer’s recommendations by a thermocycler (BIO-RAD CFX96™ Real-Time System). The primers for RT-qPCR targeted the N protein (S segment) of EBIV, including EBIV-NP-F (5’-GGTACCTCTGGCGCATTGTCTTTTC-3’), EBIV-NP-R (5’-GAAAAATGGCATCACCTGGGAAAGT-3’), and EBIV-NP-Probe (5’-FAM-TTTTGGGTCCATCTCTTTCCTCTGC-BHQ1-3’). Both of primers were synthesized by TSINGKE (Wuhan Branch, China). Twenty microliter reaction mixtures containing 2 μL of viral RNA and 0.8 μL of each primer were incubated at 55□ for 10min and 95□ for 1 min followed by 40 cycles of 95□ for 10 s and 55□ for 30 s.

### 2.8 Cell tropism of rEBIV/eGFP/S

Cells from different sources, including Vero E6, SW13, PK15, LMH and C6/36 cells, were seeded in 6-well plates, respectively. After incubated overnight the cells were infected with the rEBIV/eGFP/S virus at MOI of 0.1. After 1 h incubation in 37 °C, the virus diluent was discarded, and fresh cell culture medium was added then cultured at 37 °C. The cells would be observed and taken the pictures were taken every day (from day 0 to day 7) to record the virus infection.

### 2.9 High-content Screening assay conditions

The cell density, infective dose, and assay endpoint were the same as the previous described (27). Cell densities (10,000 cells per well) of BHK-21 cells were infected at MOI values 0.01. And two drugs Ribavirin and Favipiravir were serially diluted ranging from 50 μM to 0.78125 μM to obtain the EC50. Fluorescent signal was detected by Operetta imaging system (Perkin Elmer) at 36 h after rEBIV/eGFP/S inoculation, and the cell viability was detected under a microscope at the same time. Statistical calculations of Z’ - values were made as follows: Z’ = 1 – (3SD of sample + 3SD of control) / (|Mean of sample – Mean of control|) (28). Here, SD is the standard deviation of the fluorescent signals from cell control or sample. Z’ value is meaningful within the range of −1 < Z’≤1, the larger the value the higher the data quality, and between 0.5 and 1 are considered good quality. Then the experiment was repeated on EBIV-WT by plaque assay to confirm the antiviral results.

A library of 96 compounds from natural extracts was purchased from Weikeqi Biotech (Sichuan, China). Compounds were stored as 20□mM stock solutions in DMSO at –80°C until use. BHK-21 cells were dissociated and seeded at a density of 1□×□10^4^ cells per well in 96-well plates. After overnight incubation, cell monolayers were treated with the compounds at a final concentration of 10□μM and at the same time infected with the rEBIV/eGFP/S at the MOI of 0.01. DMEM with 2% FBS and 0.1% DMSO was negative control, and 50 μM Ribavirin was used as positive control. After 36 h incubation, the fluorescent signal in all wells was detected by Operetta imaging system (PerkinElmer) and the Z’ factor was calculated and analyzed.

## 3 Result

### 3.1 Construction and characterization of rEBIV/WT and rEBIV/eGFP/S

We constructed an infectious cDNA clone of EBIV which was isolated from *Culex modestus* mosquitoes in 2014 in Xinjiang (20). As depicted in Fig. 1A, C, D, three segments covering the complete genome sequence of EBIV were chemically synthesized and then cloned into a low-copy-number vector as described in materials and methods. We also designed the EBIV reporter virus with eGFP gene using the similar strategy used for CCHFV/ZsG reporter virus (29). The eGFP gene with P2A was inserted between the 5’UTR and N protein-ORF region of pLCK-EBIV-S (Fig. 1B). The resulted clone was designated as pLCK-EBIV-eGFP/S as described in materials and methods.

The two group of transcription plasmids (one is pLCK-EBIV-S + pLCK-EBIV-M + pLCK-EBIV-L, while another is pLCK-EBIV-eGFP/S + pLCK-EBIV-M + pLCK-EBIV-L, as shown in Fig. 1E) were transfected into BSR-T7 cells, respectively, to test the viral rescue function of the infectious clone. The supernatants were collected every day (called rEBIV/WT and rEBIV/eGFP/S) and subjected to plaque assay to determine the plaque morphology. EBIV-WT, rEBIV/WT and rEBIV/eGFP/S were seeded in BHK-21 cells respectively at MOI=0.01 for virus production, one plate with three viruses were fixed at 24 hpt (hour post transfection) and subjected to IFA using specific antibody against EBIV N protein to detect viral protein synthesis. The supernatants form another plate with three viruses were collected every day and their titer were determined to study the growth dynamics of three viruses.

As shown in Fig. 2A and 2B, when transfected the transcription plasmids for rEBIV, the CPE appeared at 24 hpt and got obvious at 48 hpt. While for rEBIV/eGFP/S, although the bright green fluorescence could be observed on the second day of transfection, the CPE appeared on the 4 dpt, and got apparent on 5 dpt (SFig 1). The rescue efficiency could get to 70% (positive detection in 7 of 10 replica wells) in once successive rescue experiment (results not shown). The IFA results showed (Fig. 2C) that the IFA-positive cells of rEBIV/WT were almost 100% among the infected cells at 24 hpt, which was similar to EBIV-WT, while the rEBIV/eGFP/S showed less IFA-positive cells at 24 hpt.

**Fig 2.**
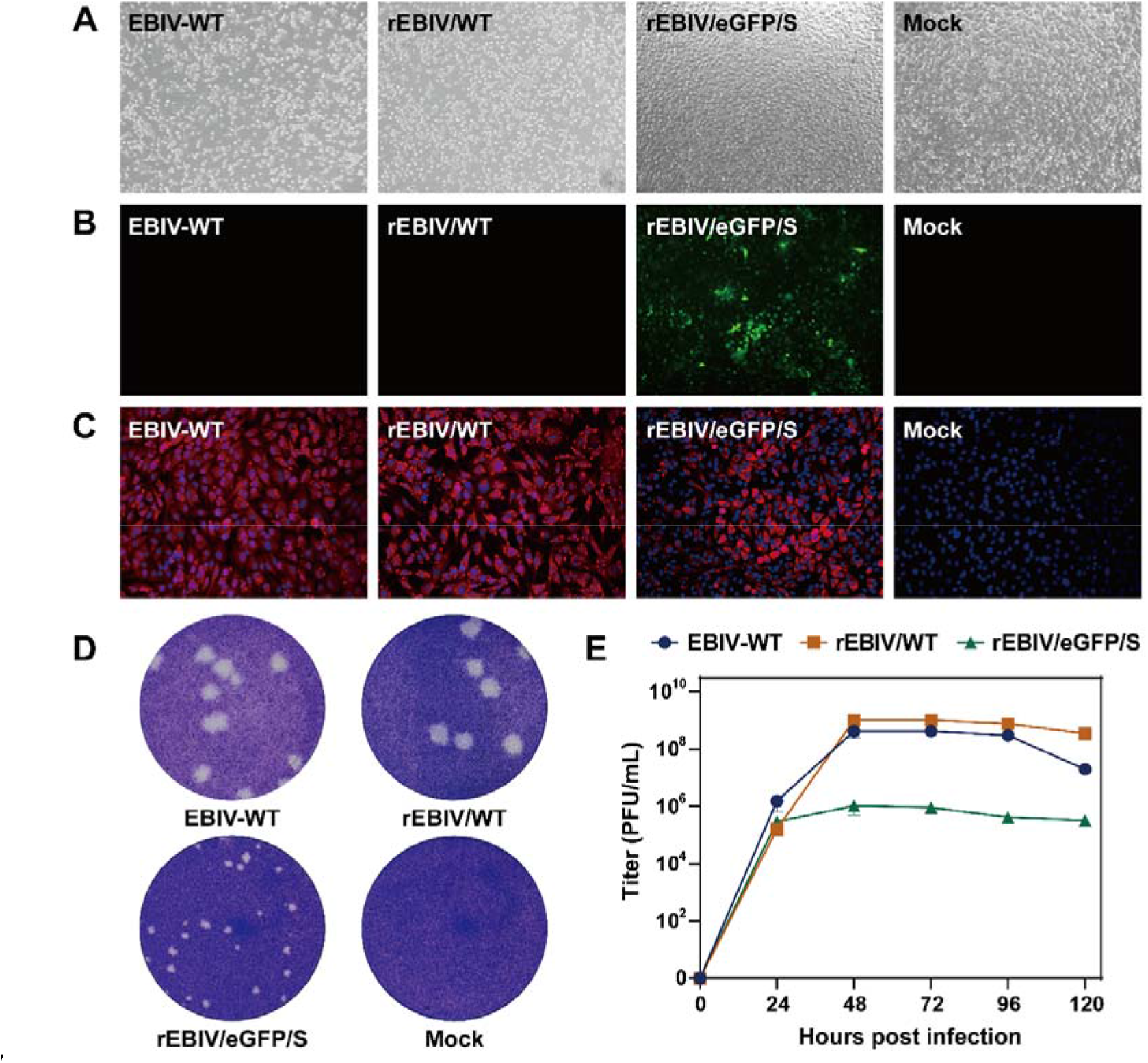
Characterization of wild type EBIV, rEBIV/WT and rEBIV/eGFP/S. (A) the cell state at 48 hpt. (B) Analysis of eGFP expression in the BSR-T7 cells and the expression of eGFP was detected under a fluorescent microscope at the 48 hpt. (C) IFA of viral protein expression in BHK-21 cells infected with the three viruses. IFA was performed at the 24 hpi using the antibody against the N protein. (D) Plaque morphology of the three viruses. (E) Growth kinetics curves of wild type EBIV, rEBIV/WT and rEBIV/eGFP/S.

The viral plaque morphology for rEBIV/WT measured at different time points were homogeneously large in BHK-21 cells, besides, both shape and size seemed similar to that of wild type virus. On the other hand, the rEBIV/eGFP/S displayed obviously smaller plaques than EBIV-WT and rEBIV (Fig. 2D). At the same time, the viral growth kinetics (Fig. 2E) showed the rEBIV/WT exhibited indistinguishable patterns of replication with wild type virus in BHK-21, whereas the viral productions of rEBIV/eGFP/S were about 100-fold lower than that of wild type virus at the same MOI.

These results demonstrated that these two rescued viruses both recovered by the infectious clone replicated efficiently. However, differences in IFA positive rate, viral plaques morphology and viral kinetics between EBIV-WT and rEBIV/eGFP/S, illustrate that the insertion of the eGFP reporter gene into EBIV genome affected the viral replication in BHK-21 cell.

### 3.2 Stability of the rEBIV/eGFP/S virus in cell culture

To analyze whether the eGFP reporter gene can be stably maintained in cell culture, the rEBIV/eGFP/S virus was serially passaged in BHK-21 cells for ten rounds. As shown in Fig. 3A, all of the BHK-21 cells infected by P1-P10 reporter viruses showed strong fluorescence signals and nearly 100% were eGFP positive when the apparent CPE appeared, indicating the eGFP gene was stably maintained during passaging. In addition, for each passage, the RNAs of the infected cells were extracted and subjected to RT-PCR to test gene stability. Different sizes of bands were expectedly detected for WT (151 bp) and reporter virus (868 bp) as the insertion of the eGFP gene (Fig. 4B). Each of the P1-P10 RNAs extracted from BHK-21 cells displayed a specific band showing no sequence deletion within the reporter gene which is confirmed by sequencing, further suggesting the stability of the reporter virus in BHK-21 cells.

**Fig 3.**
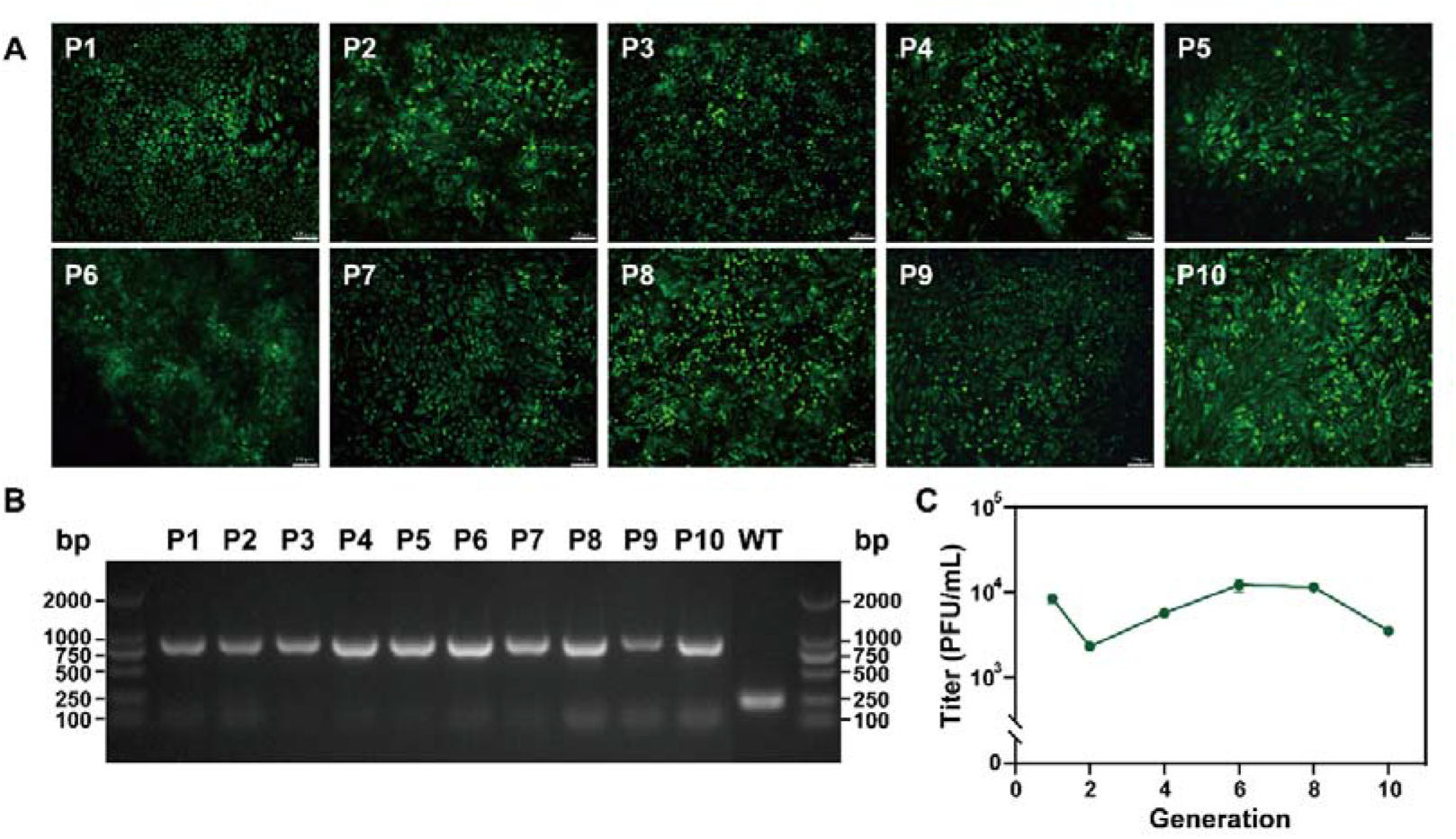
Genetic stability of rEBIV/eGFP/S in BHK-21 cells. (A) The eGFP expression of the different passages of rEBIV/eGFP/S in BHK-21 cell. The rEBIV/eGFP/S was serially passaged in BHK-21 cells for ten rounds. The expression of eGFP was detected under a fluorescent microscope at 48 h after infection. (B) Detection of the eGFP gene during virus passage in BHK-21 cell. Total RNAs from the infected cells were extracted and subjected to RT-PCR detection using the primers spanning 5’UTR to N protein gene that include the complete eGFP gene. The resulting RT-PCR products were resolved by 1% agarose gel electrophoresis. (C) The titer of 1^st^ to 10^th^ generation of rEBIV/eGFP/S. The virus titers were calculated by RT-qPCR results.

**Fig. 4.**
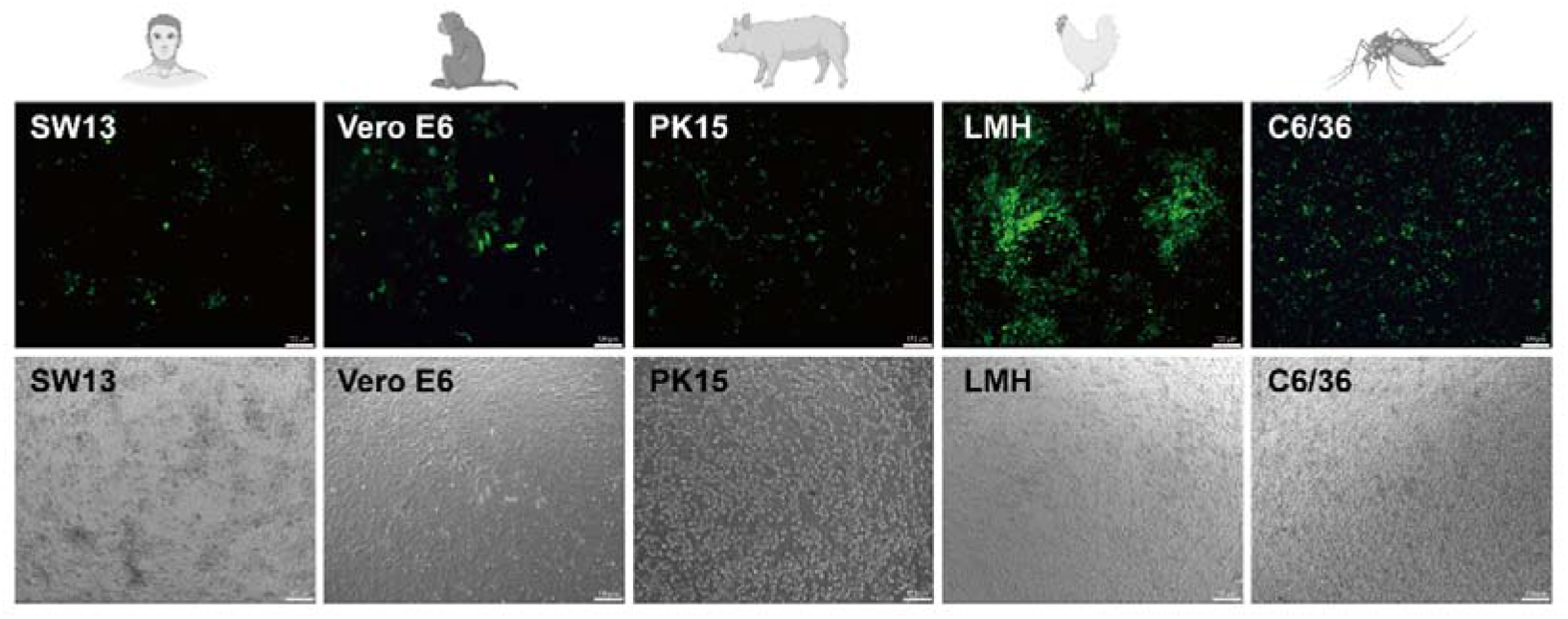
Cell lines derived from human, monkey, pig, avain, and mosquito infected with rEBIV/eGFP/S.

### 3.3 Wide cell tropism and efficient replication in different cell cultures

To further confirm the viral infectivity to cells of different origin, five cell lines were selected and inoculated with the rEBIV/eGFP/S at MOI=0.1. Pictures were taken from D1 to D10 to observe fluorescence in cell. As shown in Fig 4, all five cell lines could be infected by rEBIV/eGFP/S, but with different sensitivities. LMH cells were highly susceptible to rEBIV/eGFP/S, suggesting that EBIV could be transmitted among avian species, which consistent with previous reports (22). Besides, the rEBIV/eGFP/S is also high infectious to C6/36 cells, just as we speculated, since EBIV is an arbovirus that isolated from mosquitoes. However, SW13, Vero E6 and PK15 showed a bit lower susceptible to rEBIV/eGFP/S when compared to LMH and C6/36 cells.

### 3.4 Antiviral activity evaluation based on the rEBIV/eGFP/S reporter virus

Ribavirin and favipiravir (also known as T-705) were both broad-spectrum antiviral drugs that found to inhibit the RdRp of RNA viruses (30)(31), and had been reported to have inhibitory effect on several orthobunyaviruses replication, such as LACV (32)(33), BUNV (34), Jamestown Canyon virus (JCV) (33). In order to validate the utility of rEBIV/eGFP/S reporter virus for antiviral screening, we compared the antiviral ability of ribavirin and favipiravir on rEBIV/eGFP/S. BHK-21 cells were infected with rEBIV/eGFP/S at an MOI of 0.01 and respectively treated with different final concentrations of ribavirin and favipiravir (0 μM-50 μM) at the same time. At 36 hpi, the fluorescence value was read by Operetta imaging system (PerkinElmer), and the cell viability was detected under a microscope. The inhibition rate was derived from the ratio of the fluorescence value of the concentration to the negative contrast from the fluorescence number of each concentration minus the negative contrast, and divided by the negative contrast. The inhibition rate in the rEBIV/eGFP/S infected BHK-21 increased dramatically in a dose-dependent manner of ribavirin (Fig. 5A). The EC50 of ribavirin calculated by inhibition rate was 21.91 μM. However, the favipiravir was inactive against rEBIV/eGFP/S even at 50 μM (Fig. 5B), which is similar with the wild type EBIV (Fig. 5C, D). These results indicated that the anti-EBIV activity of compound can be rapidly evaluated by eGFP signal detection of the rEBIV/eGFP/S infected cells.

**Fig. 5.**
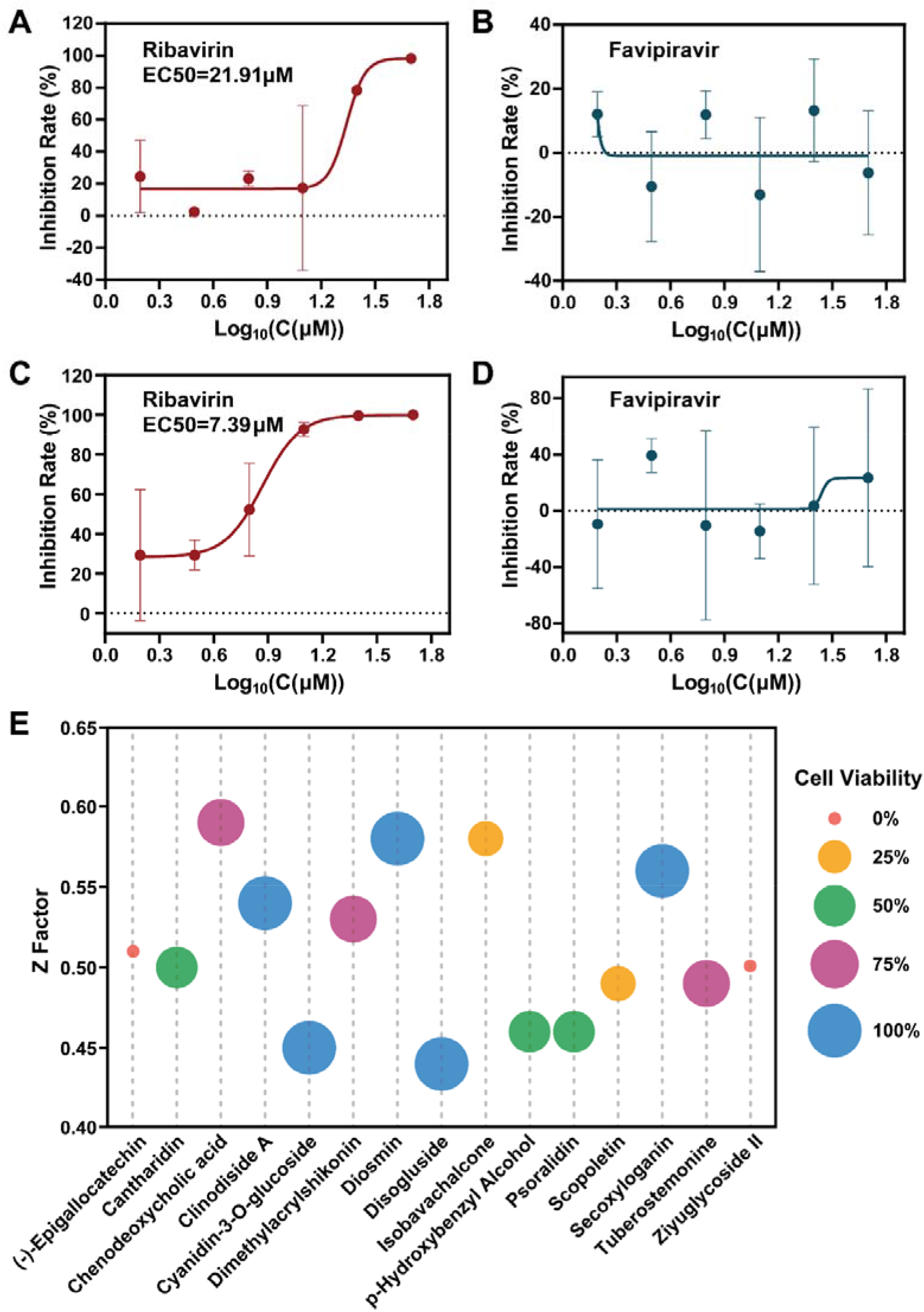
Antiviral activity of ribavirin, favipiravir and some medicines on rEBIV/eGFP/S. (A) Inhibition rate of different concentrations of ribavirin (0-50 μM) on the eGFP expression of the rEBIV/eGFP/S infected cells at 36 hpi. The EC50 was calculated by nonlinear regression using Prism software (GraphPad) as shown was 21.91 μM. Error bars indicate the standard deviations from three independent experiments. (B) Inhibition rate of rEBIV/eGFP/S infected BHK-21 cells treated with different concentrations of favipiravir. And favipiravir had no inhibitory effect on rEBIV/eGFP/S. (C), (D) Inhibition rate of different concentrations of ribavirin and favipiravir (0-50 μM) on wild type EBIV, tested by plaque assay. (E) The library of compounds was scanned, and the Z factor of each compound and positive control were obtained by the fluorescence value. The compounds that had higher Z factor than positive controls were shown in the table. The different colors and sizes of circles indicated their respective cell viability.

We further chose 25 μM ribavirin as positive control, and tested the anti-rEBIV/eGFP/S activity of the library of compounds from nature product. The BHK-21 cells were infected with rEBIV/eGFP/S at an MOI of 0.01 and treated with various compounds (10 μM). The fluorescence value was read at 36 hpi, and the cell viability was observed at the same time. The Z’ factor of the positive and negative control is 0.46, indicates that this model is not very good but acceptable. As depicted in Fig. 5C and several compounds showed a significant inhibitory effect on rEBIV/eGFP/S replication with higher Z factors (the raw data of the screening is in STable). The wells with Clinodiside A, Diosmin, Secoxyloganin, Disogluside, and Cyanidin-3-O-glucoside showed both high Z factors and 100% cell viability, proving they may be potential effective anti-EBIV drugs. Overall, these results demonstrated that the rEBIV/eGFP/S reporter virus provides a rapid and precise tool for antiviral inhibitors screening against EBIV.

## 4 Discusssion

EBIV as a newly classified orthobunyavirus as a new species in the *Peribunyaviridae* showed potential threat to human or animal health. In order to develop reliable tools for EBIV study, here we successfully constructed the infectious clones of EBIV and an eGFP reporter virus rEBIV/eGFP/S. By the standard virus rescue procedure, we identified that the rEBIV/WT virus showed indistinguishable replication efficiency with the wild type EBIV (Cu20-XJ) in BHK-21 cells, while the insertion of eGFP reduced the replication efficiency. The eGFP gene expression level within the rEBIV/eGFP/S infected cells correlated well with the viral replication, inferring that the growth of reporter virus can be monitored directly by eGFP observation. The rEBIV/eGFP/S could stably passage in BHK-21 cells and show different tropism on cell lines of different sources. After validating the same medicines (ribavirin and favipiravir) had the same inhibitory effect on both wild type EBIV and rEBIV/eGFP/S in BHK-21, we confirmed the feasibility of the rEBIV/eGFP/S for rapid antiviral screening assay, and five compounds were found to have an inhibitory effect on the EBIV.

In our study, the eGFP gene was first inserted into the N or C-terminal of N protein directly, however, neither approach rescued the virus. We considered that this may because the eGFP affected the normal expression and native structure of N protein, leading to the failure of the virus to replicate and package normally. Then we added P2A between eGFP and N protein, allowing separate expression of these two proteins, to get infectious rEBIV/eGFP/S. But the results of plaque assays and growth curves suggested that the virus was significantly different from the wild type, possibly because of the insertion of eGFP may affect the expression of NSs protein, which has multiple functions in the viral replication cycle and is the major virulence factor (35) but dispensable for virus growth (36)(37). We also tried to insert the P2A with eGFP between 3’ UTR and N protein ORF, but also failed. The reason would be when the N protein precedes P2A, it will carry more than 20 amino acid tails during translation shearing, which may affect the function of N protein. As for other two segments, there has been successful rescued cases on M segment in Bunyamwera virus (38), which replaced almost half of the N terminal of Gc to eGFP. However, insert report gene to L segment has not been studied yet, which may due to the large size of L and potential difficulty to modify the RdRp and preserving its polymerase activity.

To evaluate the rEBIV/eGFP/S stability in both mammalian and mosquito cell lines, we also did continuous passages of rEBIV/eGFP/S on C6/36 cells. To our surprise, although the fluorescence could be observed and the eGFP gene could be detected by RT-PCR using specific eGFP primers, the eGFP gene seemed to switch places since the bands significantly shorter when using the primers for sequences outside the eGFP from the second generation (SFig 2). The similar gene loss with unknown mechanism has been reported in other viruses in C6/36 cells(39)(40)(41), we speculated that due to the different replication mechanisms of EBIV between mammalian and insect cells, the insertion of an additional ORF into the viral genome may affect RNA replication and the stability of inserted gene in C6/36 but not in BHK-21 cells. Further work is needed to test this hypothesis.

Ribavirin is the first synthetic nucleoside analogue that has ever been reported to be active against a broad spectrum of RNA viruses (such as hepatitis C virus (HCV), Respiratory Syncytial Virus (RSV), and influenza virus) (42). And it can also reduce the replication of EBIV, and the EC50 of rEBIV/eGFP/S is 21.91 μM which is similar to BUNV-mCherry (34). Favipiravir triphosphate shows broad-spectrum inhibitory activities against the RNA polymerases of influenza A viruses (including the highly pathogenic H5N1 viruses) (43) and many other positive-sense RNA (such as West Nile virus (WNV) and Western equine encephalitis (WEE)) and negative-sense RNA viruses (such as Crimean-Congo hemorrhagic fever virus (CCHFV), Severe fever with thrombocytopenia syndrome virus (SFTSV), Rift valley fever virus (RVFV), and Ebola virus) (31). But in our study, the favipiravir is ineffective even at higher concentrations, the specific reasons need to be further explored. As for the antiviral candidates we found in this research, diosmin and cyanidin-3-O-glucoside were identified as inhibitor of SARS-CoV-2, since diosmin could bind covalently to the SARS-CoV-2 main protease, inhibiting the infection pathway of SARS-CoV-2 (44), and cyanidin-3-O-glucoside was demonstrated to inhibit M protein activity of SARS-CoV-2 in a dose-dependent manner at biologically relevant (μM) concentrations (45). The other three compounds, clinodiside A, secoxyloganin and disogluside, have not been reported to have antiviral effects yet. More antiviral mechanisms and minimum effective dose need to be further studied.

In conclusion, we have established the reverse genetic system for EBIV, and rescued a reporter virus rEBIV/eGFP/S. The reporter virus showed good stability in BHK-21 cell and different tropism in various cell lines. The reporter virus based antiviral assay developed in this study will facilitate the antiviral screening for novel anti-EBIV agents.

## Supporting information

SFig.1

SFig. 2

STable

## Author Contributions

HX and ZY designed the experiments. NR, FW, LZ, LW, and JQ performed the experiments. NR, FW, and JQ analyzed the data. GZ, EB, and BZ contributed the reagents, materials, and analysis tools. NR, FW, EB, ZY and HX wrote and review the manuscript. All authors contributed to the article and approved the submitted version.

## Acknowledgments

We would like to thank the valuable suggestions from Prof. Ke Peng (Wuhan Institute of Virology) and Dr. Shufen Li (Wuhan Institute of Virology) to our experiment, and Ding Gao and An-Na Du from the Core Facility and Technical Support, Wuhan Institute of Virology, for their help with HCS.

