## Supplementary figures and images for "Efficient rescue of a newly classified Ebinur lake orthobunyavirus with GFP reporter and its application in rapid antiviral screening"

### SFig.1

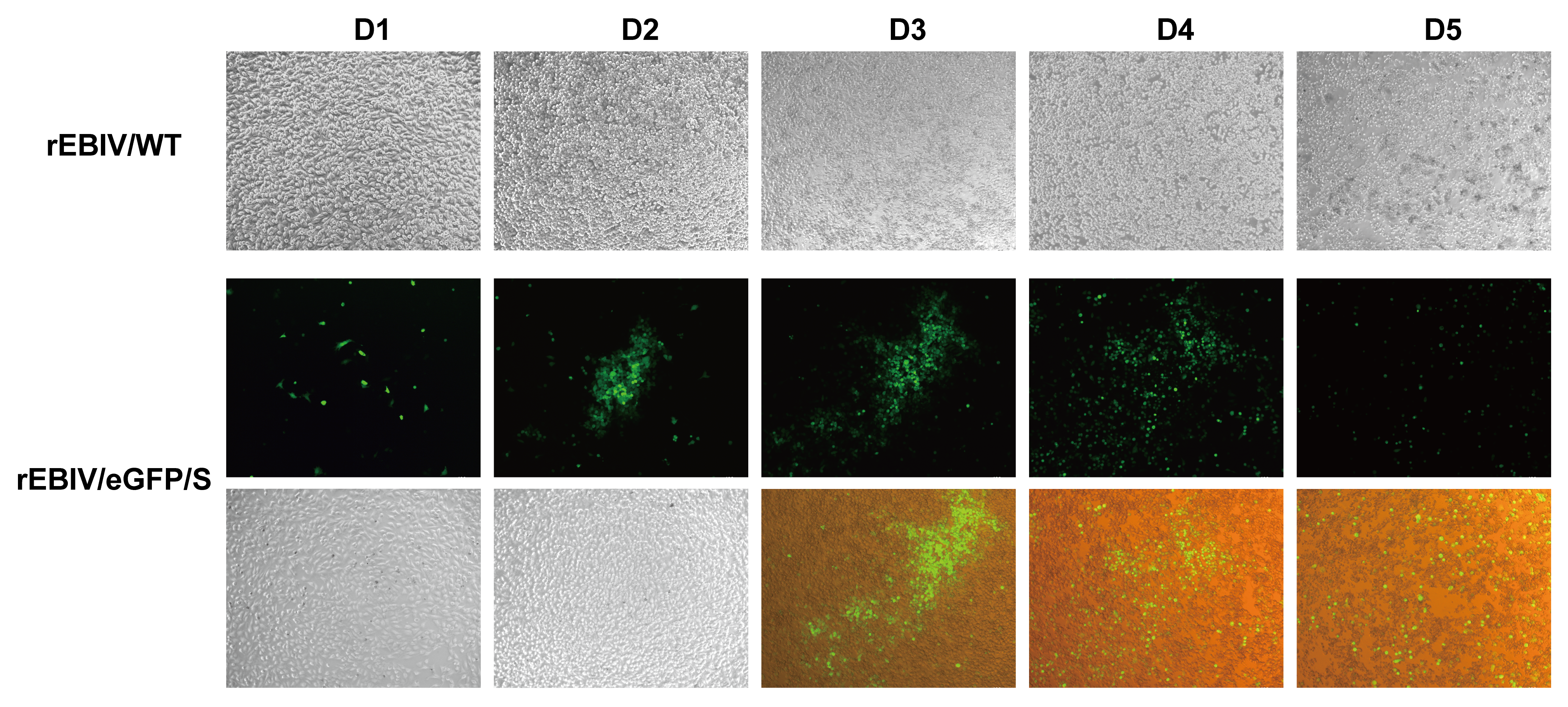

### SFig. 2

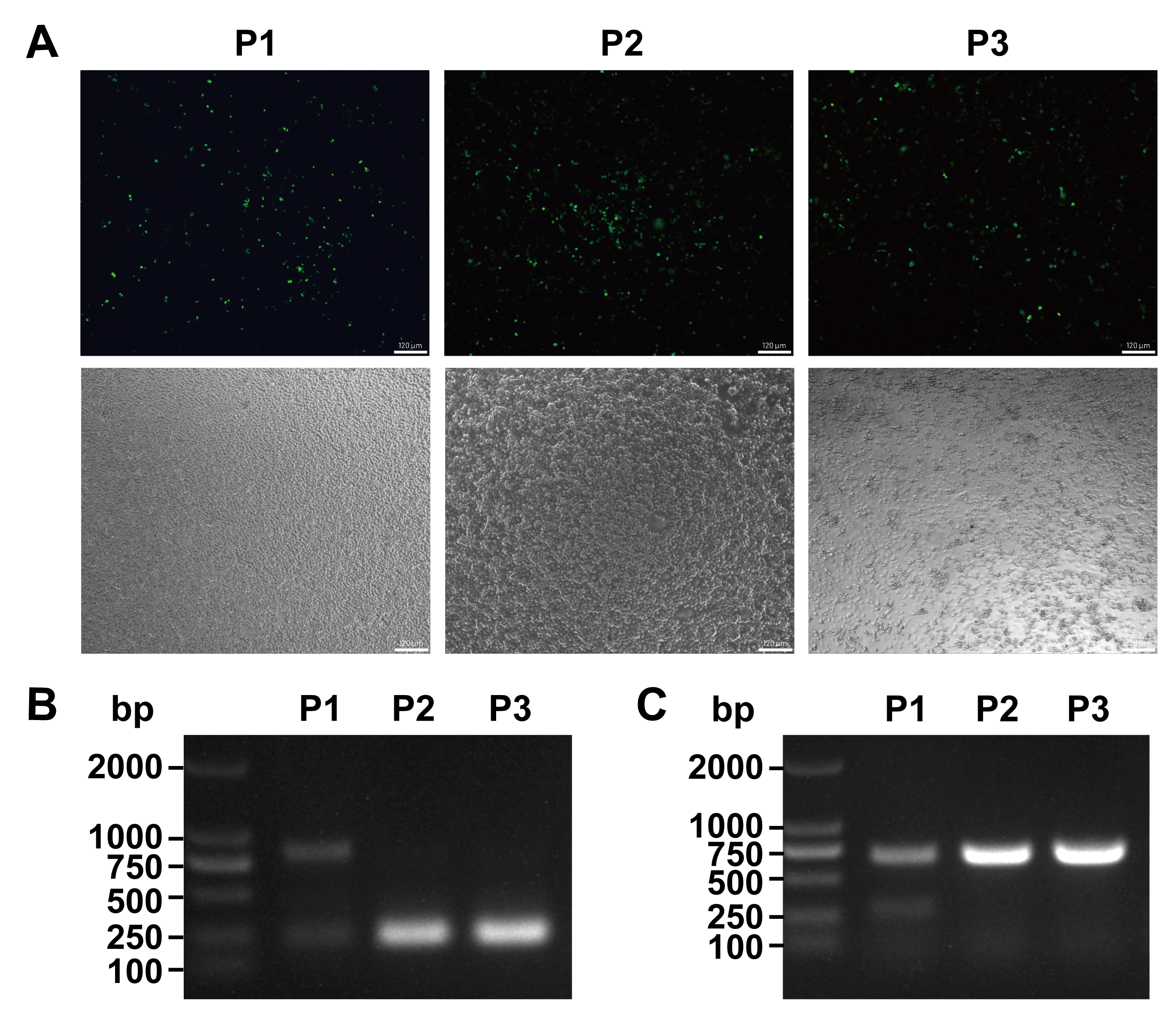
