## Supplementary material for "Efficient rescue of a newly classified Ebinur lake orthobunyavirus with GFP reporter and its application in rapid antiviral screening": STable

**STable: The results of high-content screening**

| **Number** | **Name** | **Number of eGFP** | | **Z Factor** |
| --- | --- | --- | --- | --- |
| 1 | Psoralen | | 1639 | -3.66 |
| 2 | Psoralidin | | 369 | 0.46 |
| 3 | Bakuchiol | | 955 | -0.11 |
| 4 | Bavachin | | 1482 | -23.05 |
| 5 | Isobavachalcone | | 33 | 0.58 |
| 6 | Epicatechin | | 1223 | -1.16 |
| 7 | (-)-Epigallocatechin | | 255 | 0.51 |
| 8 | Helicid | | 492 | 0.40 |
| 9 | Peiminine | | 1170 | -0.82 |
| 10 | Peimine | | 740 | 0.20 |
| 11 | Peimisine | | 852 | 0.06 |
| 12 | Brazilin | | 611 | 0.32 |
| 13 | Scoparone | | 1502 | -110.50 |
| 14 | Menthol | | 564 | 0.35 |
| 15 | Cyasterone | | 564 | 0.35 |
| 16 | Stigmasterol | | 632 | 0.30 |
| 17 | Cantharidin | | 286 | 0.50 |
| 18 | D-Galactose | | 1256 | -1.44 |
| 19 | 10-Deacetylbaccatin Ⅲ | | 1186 | -0.91 |
| 20 | Apigenin | | 1269 | -1.57 |
| 21 | Mandelic acid | | 1041 | -0.31 |
| 22 | Bicucullin | | 1357 | -3.07 |
| 23 | Menthone | | 826 | 0.10 |
| **Number** | **Name** | **Number of eGFP** | | **Z Factor** |
| 24 | Cardamonin | | 936 | -0.07 |
| 25 | Kaurenoic acid | | 1393 | -4.36 |
| 26 | Palmatine chloride | | 1329 | -2.44 |
| 27 | Phenyl benzoate | | 1620 | -4.45 |
| 28 | Curcumol | | 828 | 0.10 |
| 29 | Curdione | | 1037 | -0.30 |
| 30 | Furanodienon | | 806 | 0.13 |
| 31 | Curcumenol | | 944 | -0.09 |
| 32 | Farrerol | | 1651 | -3.27 |
| 33 | Chenodeoxycholic acid | | 11 | 0.59 |
| 34 | Catechin | | 1029 | -0.28 |
| 35 | 1,8-Dihydroxyanthraquinone | | 1573 | -8.36 |
| 36 | 2,3,5,4＇-Tetrahydroxy stilbene-2-Ο-β-D-glucoside | | 449 | 0.42 |
| 37 | 1,3-Dicaffeoylquinic acid | | 814 | 0.12 |
| 38 | Dimethylacrylshikonin | | 213 | 0.53 |
| 39 | 3,6'-Disinapoyl sucrose | | 759 | 0.18 |
| 40 | Clinodiside A | | 183 | 0.54 |
| 41 | Ligustroflavone | | 1284 | -1.74 |
| 42 | Tannic acid | | 1253 | -1.41 |
| 43 | Cholic acid | | 849 | 0.07 |
| 44 | Angelic Acid | | 1151 | -0.72 |
| 45 | Mangostin | | 1521 | -44.43 |
| 46 | Scutellarin methyl ester | | 1230 | -1.21 |
| 47 | Diosmin | | 59 | 0.58 |
| **Number** | **Name** | **Number of eGFP** | | **Z Factor** |
| 48 | Secoxyloganin | | 114 | 0.56 |
| 49 | Eugenol | | 335 | 0.37 |
| 50 | Syringaresnol-4-O-β-D-apiofuranosy | | 511 | 0.27 |
| 51 | N-Butylscopolammonium Bromide | | 520 | 0.26 |
| 52 | Syringate | | 1993 | -1.11 |
| 53 | Scopolin | | 1501 | -6.34 |
| 54 | Scopolamine | | 743 | 0.07 |
| 55 | Scopoletin | | 25 | 0.49 |
| 56 | Huperzine-A | | 474 | 0.29 |
| 57 | Ethyl 4-methylcinnamate | | 680 | 0.13 |
| 58 | p-Hydroxybenzyl Alcohol | | 118 | 0.46 |
| 59 | p-Hydroxybenzaldehyde | | 494 | 0.28 |
| 60 | p-Hydroxybenzoic acid | | 702 | 0.11 |
| 61 | p-Hydroxy-cinnamic acid | | 297 | 0.39 |
| 62 | p-Coumaric acid | | 581 | 0.22 |
| 63 | Tuberostemonine | | 22 | 0.49 |
| 64 | Complanatoside A | | 1822 | -2.82 |
| 65 | Docetaxel | | 344 | 0.36 |
| 66 | Disogluside | | 181 | 0.44 |
| 67 | Ziyuglycoside II | | 1 | 0.50 |
| 68 | Albiflorin | | 899 | -0.13 |
| 69 | Paeoniflorin | | 1571 | -19.24 |
| 70 | Tectoridin | | 1347 | -2.06 |
| 71 | Cyanidin-3-O-glucoside | | 145 | 0.45 |
| **Number** | **Name** | **Number of eGFP** | | **Z Factor** |
| 72 | Strychnine | | 366 | 0.35 |
| 73 | Dencichine | | 459 | 0.30 |
| 74 | Cephalomannine | | 709 | 0.11 |
| 75 | Harringtonine | | 725 | 0.09 |
| 76 | Umbelliferone | | 1116 | -0.63 |
| 77 | Carnosol | | 317 | 0.38 |
| 78 | L-rhamnose monohydrate | | 771 | 0.04 |
| 79 | Amentoflavone | | 593 | 0.21 |
| 80 | Dihydroartemisinin | | 1059 | -0.46 |
| 81 | Phytolaccagenin | | 826 | -0.03 |
| 82 | Spinosin | | 763 | 0.05 |
| 83 | Osthole | | 1104 | -0.59 |
| 84 | Muscone | | 1184 | -0.89 |
| 85 | Thymol | | 1286 | -1.48 |
| 86 | Cimicidanol 3-arabinoside | | 911 | -0.15 |
| 87 | Cimifugin | | 935 | -0.19 |
| 88 | Jujuboside A | | 1007 | -0.34 |
| 89 | Prim-O-glucosylcimifugin | | 874 | -0.09 |
| 90 | Eriocitrin | | 789 | 0.02 |
| 91 | Dendrobine | | 695 | 0.12 |
| 92 | Gypsogenin-3-O-β-D-gluco pyranoside | | 1347 | -2.06 |
| 93 | (S)-Tetrahydrocolumbamine | | 1073 | -0.50 |
| 94 | Berberrubine | | 672 | 0.14 |
| 95 | Berberine | | 1214 | -1.03 |
| **Number** | **Name** | **Number of eGFP** | | **Z Factor** |
| 96 | Jujuboside B | | 1578 | -23.56 |
